# Large herbivorous wildlife and livestock differentially influence the relative importance of different sources of carbon for riverine food webs

**DOI:** 10.1101/2021.07.29.454315

**Authors:** Frank O. Masese, Thomas Fuss, Lukas Thuile Bistarelli, Caroline Buchen-Tschiskale, Gabriel Singer

## Abstract

In many regions around the world, large populations of native wildlife have declined or been replaced by livestock grazing areas and farmlands, with consequences on terrestrial-aquatic ecosystems connectivity and trophic resources supporting food webs in aquatic ecosystems. The river continuum concept (RCC) and the riverine productivity model (RPM) predict a shift of carbon supplying aquatic food webs along the river: from terrestrial inputs in low-order streams to autochthonous production in mid-sized rivers. Here, we studied the influence of replacing large wildlife (mainly hippos) with livestock on the relative importance of C3 vegetation, C4 grasses and periphyton on macroinvertebrates in the Mara River, which is an African montane-savanna river known to receive large subsidy fluxes of terrestrial carbon and nutrients mediated by LMH, both wildlife and livestock. Using stable carbon (δ^13^C) and nitrogen (δ^15^N) isotopes, we identified spatial patterns of the relative importance of allochthonous carbon from C3 and C4 plants (woody vegetation and grasses, respectively) and autochthonous carbon from periphyton for macroinvertebrates at various sites of the Mara River and its tributaries. Potential organic carbon sources and invertebrates were sampled at 80 sites spanning stream orders 1 to 7, various catchment land uses (forest, agriculture and grasslands) and different loading rates of organic matter and nutrients by LMH (livestock and wildlife, i.e., hippopotamus). The importance of different sources of carbon along the river did not follow predictions of RCC and RPM. First, the importance of C3 and C4 carbon was not related to river order or location along the fluvial continuum but to the loading of organic matter (dung) by both wildlife and livestock. Notably, C4 carbon was important for macroinvertebrates even in large river sections inhabited by hippos. Second, even in small 1^st^ −3^rd^ order forested streams, autochthonous carbon was a major source of energy for macroinvertebrates, and this was fostered by livestock inputs fuelling aquatic primary production throughout the river network. Importantly, our results show that replacing wildlife (hippos) with livestock shifts river systems towards greater reliance on autochthonous carbon through an algae-grazer pathway as opposed to reliance on allochthonous inputs of C4 carbon through a detrital pathway.

## 1. Introduction

While resource transfers from terrestrial to riverine aquatic ecosystems can occur passively through wind and atmospheric deposition, overland and riverine flow (Abrantes and Sheaves, 2010; Lamberti et al., 2010; Wipfli et al., 2007), animals are major active resource vectors through defecation (egestion), urination (excretion) and in the form of carcasses (Naiman and Rogers, 1997; Subalusky et al., 2015; Subalusky et al., 2017). Resource subsidies by large mammalian herbivores (LMH) are particularly interesting because of their capacity to transfer large amounts of high-quality, nutrient-rich resources that strongly influence abiotic conditions, productivity, community structure and trophic interactions in recipient aquatic ecosystems (Vanni, 2002; Masese et al., 2018; Stears et al., 2018).

Along their longitudinal gradient, rivers are postulated to exhibit a prominent shift in trophic resources supporting aquatic communities, influencing their structure and functional organization besides physical and chemical factors (Vannote et al., 1980; Atkinson et al., 2017). In temperate headwater streams, allochthonous carbon from C3-plants (woody vegetation) dominates the base of food webs. As streams widen, however, in-stream primary production grows, while allochthonous inputs decrease to subsidies leaking from upstream (Vannote et al., 1980; Thorp and Delong, 1994). In contrast, in Afrotropical savanna rivers, mega-herbivores including hippos (Hippopotamus amphibius), elephants, buffalo, other ungulates, such as wildebeest, transfer substantial amounts of terrigenous carbon and nutrients to entire river networks (Naiman and Rogers, 1997; Mosepele et al., 2009, Hulot et al., 2019), challenging existing models of riverine ecosystem functioning. Notably, this allochthonous carbon is mainly derived from C4 plants in surrounding grasslands and seems to critically need grazing LMH as subsidy vectors (Marwick et al., 2014). Indeed, hippo egestion can account for up to 88% of total carbon inputs into lowland rivers (Subalusky et al., 2015), which may play a similar role as leaf litter in headwater streams in stimulating microbial processes and supporting food webs (Webster, Benfield et al. 1999). Large quantities of hippo dung have also been found to be detrimental to aquatic biodiversity, with fish kills occurring during phases of reduced flow and high hippo density (Dutton et al., 2018a; Stears et al., 2018).

Populations of native LMH have declined in their natural range and have been replaced by human settlements, farmlands and livestock (Doughty et al., 2013; Ogutu et al., 2016; Veldhuis et al., 2019). Large herds of livestock (mainly cattle, goats and sheep) may compensate losses of resource subsidies to aquatic ecosystems arising from wildlife declines (Iteba et al., 2021; Masese et al., 2020), but the quality, quantity (rate), timing and location of inputs can substantially differ (Subalusky and Post, 2019), with ensuing implications for aquatic ecosystem functioning. For instance, nutrient-rich cattle dung was found to increase primary production relative to hippo dung (Masese et al., 2020). And, since high-quality algal carbon contributes higher proportions of carbon to metazoan biomass relative to woody vegetation and grasses (Thorp and Delong, 2002; Douglas et al., 2005; Lau et al., 2009), livestock may shift aquatic food webs to be more reliant on autochthonous sources of carbon. This may be facilitated by the increased transportability of cattle dung due to smaller particle sizes and faster leaching and remineralization (Masese et al., 2020), while hippo faeces contain large particles that settle on the river bottom and decompose slowly (Dawson et al., 2016; Dutton et al., 2020).

The Mara River (Kenya, Tanzania) traverses a landscape gradient, that makes it well suited to study the influence of land use change and altered LMH populations on the terrestrial-aquatic linkages achieved through resource subsidies. The forest in the upper reaches contrasts mixed small-scale and large-scale agriculture and human settlements in mid-reaches (Mati et al., 2008). The region hosts livestock (mainly cattle, goats and sheep) at varying densities, grazing in an overlapping distribution with wildlife (Lamprey and Reid 2004; Veldhuis et al., 2019). In the lower reaches, protected savanna grasslands in the Kenyan Maasai Mara National Reserve (MMNR) harbour over a million migratory herbivores (Ogutu et al., 2016), including >4000 hippos (Kanga et al., 2011). Per-capita input of organic matter and nutrients into streams and rivers by individual livestock is small compared to hippos (Sualusky et al., 2015; Masese et al., 2020) due to smaller body sizes and shorter time of presence in or near water, but livestock can appear in large numbers (Iteba et al., 2021). Also, resource subsidies of livestock and wildlife differ in quality, with likely impacts on aquatic ecosystem functioning. A good assessment of subsidies transferred to the Mara River critically needs comparative data for livestock and wildlife and how their inputs interact with river size.

Here, we investigate carbon sources fuelling aquatic food webs at various sites along the Mara River, specifically the interaction between river size and subsidy fluxes driven by livestock and large wildlife (hippos) at various densities. In the greater Mara River network, we sampled 80 sites in streams differing in catchment land use and LMH abundance. We used natural abundance stable isotope analysis (SIA) of carbon (δ^13^C) and nitrogen (δ^15^N) to distinguish three main sources of carbon (C3 vegetation, C4 grasses and periphyton) for macroinvertebrates. Contrary to predictions of the river continuum concept (RCC) and the riverine productivity model (RPM), we hypothesise that LMH have an overriding influence over stream size in determining sources of carbon for food webs in savanna rivers.

Specifically, in lower sections of the river, autochthonous production and C3 carbon should decrease in importance while C4 carbon subsidies by livestock and large wildlife should become more prominent in the diets of invertebrates.

## 2. Materials and Methods

### 2.1 Study area

This study was conducted in the Kenyan part of the Mara River catchment. Two perennial tributaries, the Nyangores and Amala rivers drain the Mau Forest, which is the most extensive tropical moist broadleaf forest in East Africa, before joining to form the Mara mainstem in the lowlands (Fig. 1). In the middle reaches, several seasonal tributaries, including the Talek, Olare Orok, Ntiakntiak, Molibany and the Sand, drain the semi-arid livestock grazing lands and wildlife conservancies outside the Maasai Mara National Reserve (MMNR). Annual rainfall varies from about 2000 mm in the highlands to around 1000 mm in the lowland savanna (Jackson and McCarter, 1994). January to March is typically a dry period, while March-July and October-November are wet periods known as long and short rains, respectively.

**Figure 1.**
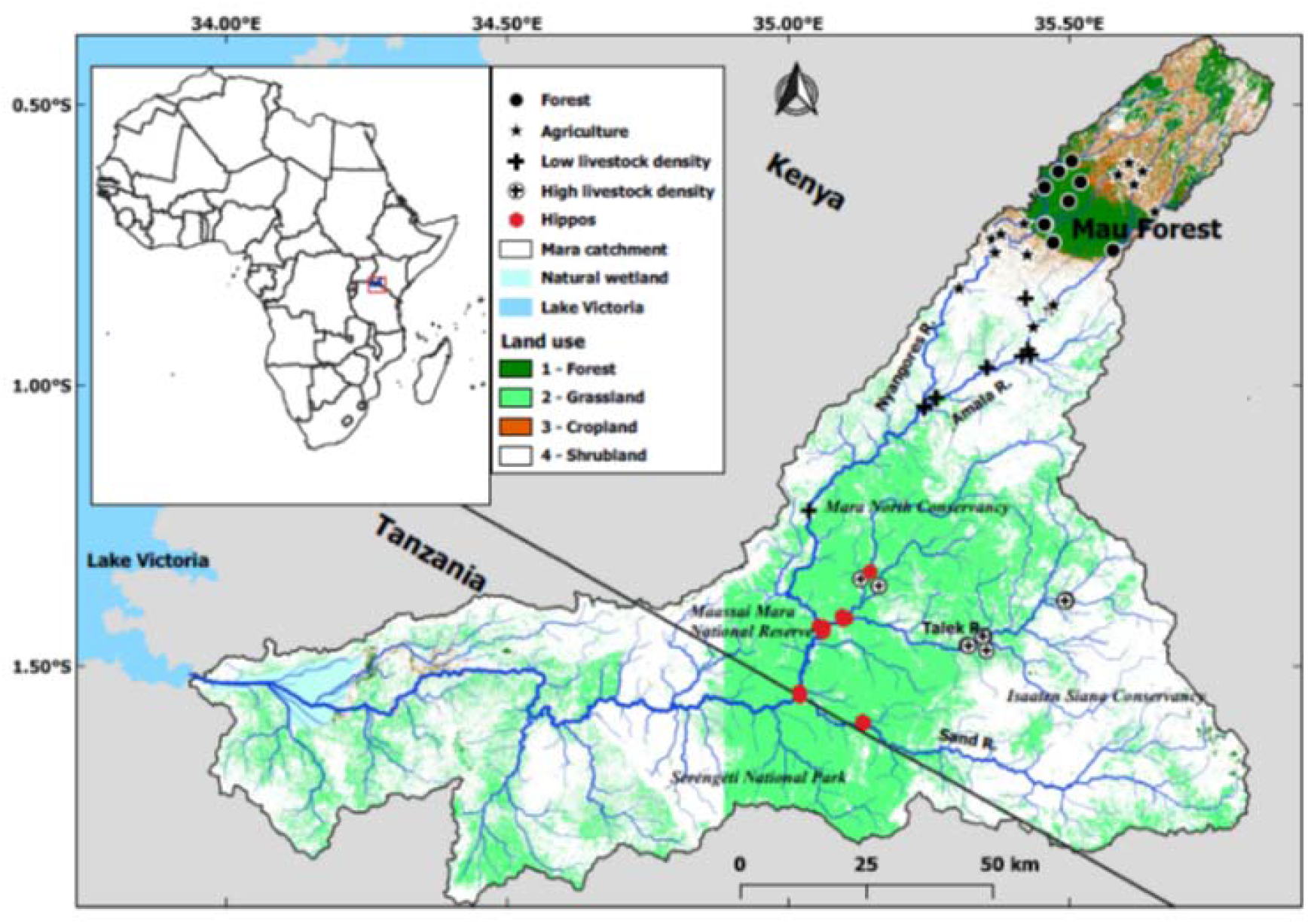
Map of the Mara River catchment showing the position of the sampling sites.

The Kalenjin ethnic group live on the highlands while the Maasai pastoralists occupy the middle and lower portions of the basin. The basin hosts substantial numbers of livestock but densities differ across the catchment as the Kalenjin have diversified to crop farming and husbandry of small herds of improved breeds of cattle, while the Maasai’s large herds of cattle are the mainstay of their livelihoods (Lamprey and Reid, 2004). Agricultural expansion is ongoing across the basin (Mati et al., 2008).

The middle reaches of the Mara River and its tributaries within the MMNR host >4,000 hippos (Kanga et al., 2011), which graze in savanna grasslands at night and rest in or near the river during the day, transferring ~36,000 kg of organic matter in the form of dung per day from the terrestrial to the aquatic domain (Kanga et al., 2011; Subalusky et al., 2015). In the same region, Maasai pastoralists graze nearly 200,000 cattle and large numbers of goats and sheep in communal lands adjoining the MMNR and utilize rivers as watering points (Lamprey and Reid, 2004; Ogutu et al., 2016; Veldhuis et al., 2019). In communal conservancies outside the MMNR, people graze their livestock in a manner that allows livestock to co-exist with wildlife (Kanga et al., 2013). This distribution results in a displacement pattern with hippo areas inside the reserve, mixed hippo and livestock (mainly cattle) areas in the conservancies and only livestock grazing areas outside the conservancies (Kanga et al., 2013).

### 2.2 Study design

A total of 80 sites were selected for sampling during the beginning of the dry season in January 2018. All sites were sampled only once unless specified otherwise. In most cases, we sampled at confluences, i.e., working in both tributaries and a third downstream mainstem site after effective mixing of water from the two tributaries. Sites were selected depending on both catchment-scale (mainly land use) and reach-scale influences by human activities, cattle and wildlife. Sites were then grouped into 5 broad regions: forested (19), agricultural (26), low-density livestock (15), high-density livestock (12), and wildlife (i.e., hippo) (10) sites. Forest sites had C3 vegetation dominating the catchment and riparian areas, and were used as a reference for the human and LMH influences. Agricultural sites were in farming areas (crop cultivation), although most farmers in the area also own low numbers (<20 per km^2^) of livestock (mainly cattle). The low-density livestock sites were also in agricultural areas but had a higher density of livestock (20-50 per km^2^). The high-density livestock sites were located in conservancies outside the MMNR where the only land use activity is the grazing of large herds of cattle (> 100 heads per km^2^), goats and sheep; although some wildlife such as zebra, wildebeest and other herbivores (but rarely hippos) also occur in these areas. Finally, hippo sites were located on the Mara River mainstem and tributaries in the MMNR downstream of river sections inhabited by large populations of hippos (Kanga et al., 2011).

### 2.3 Sample collection

At each site, samples of the dominant riparian plants were collected by hand into aluminium envelopes. Macrophytes were rare and not considered to be an important resource. We collected triplicate samples of cattle and hippo dung when encountered at a site. Triplicate periphyton samples were scrubbed from submerged surfaces after gentle washing to remove invertebrates and debris. At each site macroinvertebrates were collected from riffles, runs, pools and marginal vegetation using a dip net (500 μm mesh-size) and sorted alive into Eppendorf vials. Plant, dung, periphyton slurry (in HDPE bottles) and invertebrates were frozen for transport to the laboratory pending further processing.

#### 2.3.1 Physical and chemical variables

To explore relationships between catchment land use and the importance of various sources of carbon for invertebrates in the Mara River, sub-catchments for all sites were delineated based on a digital elevation model of Kenya (90 m resolution, Shuttle Radar Topography Mission) and their land cover analyzed according to the major types; forest (both natural and plantation), agriculture (cropland) and settlements, grasslands (grazing lands) and shrublands.

At each site, we measured average stream width at 20-30 randomly located transects and average depth and velocity at a minimum of 100 randomly located points. Discharge was estimated using the velocity-area method at one transect. Dissolved oxygen concentration (DO), pH, temperature and electrical conductivity were measured *in situ* using a WTW Multiprobe 3320 (pH320, OxiCal-SL, Cond340i; Weilheim, Germany).

### 2.3 Sample preparation and stable isotope analysis

On return to the laboratory, samples were immediately prepared for isotope analysis or preserved at −20°C. To clean periphyton, the slurry was decanted onto a petri dish and excess water evaporated in an oven at 60 °C for 48 h. The dry sample was ground using a mortar and pestle, an aliquot was weighed into a tin cup. Macroinvertebrate specimens were thawed and examined under a dissecting microscope to confirm identity and remove stomach contents before drying at 60 °C for 48 h. These samples were ground with an IKA A 11 Basic mill (IKA-Werke GmbH & Co. KG, 79219, Staufen, Germany) and weighed into tin cups.

Macroinvertebrates were identified mostly to the genus level. Several individuals of a given taxon from each site were pooled to produce sufficient dry tissue mass except for species with larger body sizes such as crabs (Potamonautes spp.), dragonflies and damselflies (Odonata), some beetles (Coleoptera) and bugs (Hemiptera). To avoid contamination by carbonates, which can be enriched in ^13^C compared to living tissue, we only used soft tissues from crabs and Mollusca.

Stable isotope analysis was done on a Thermo Finnigan DeltaPlus Advantage isotope ratio mass spectrometer (IRMS, Thermo Scientific) coupled to an ECS 4010 EA elemental analyzer (Costech Analytical Technologies). Stable isotope ratios (^13^C/^12^C and ^15^N/^14^N) are expressed in parts per mil (‰) deviations to the international standard, as defined by the equation: δ^13^C, δ^15^N = [(R_sample_/R_references_) - 1] × 10^3^, where R = ^13^C/^12^C for carbon and ^15^N/^14^N for nitrogen. δ^13^C values were normalized to the international scale Vienna Pee Dee Belemnite (VPDB) and δ^15^N values to atmospheric nitrogen, by analyzes of the international standards USGS40 and USGS41 (L-glutamic acid) within the sequence (Qi et al. 2003). In addition, a laboratory standard (δ^13^C = −27.11 ‰, C% = 48.12; δ^15^N = 0.48 ‰, N% = 2.17), which was calibrated directly against USGS40 and USGS41, was measured at different positions within the run. Precision, defined as the standard deviation (±1σ) of the laboratory control standard along the run was better than ±0.79‰ for C and ± 0.55‰ for N.

### 2.4 Functional feeding groups and trophic groups

Macroinvertebrate taxa were assigned to functional feeding groups (FFGs) and trophic groups based on (Masese et al., 2014) and references therein. FFGs considered included shredders feeding on coarse particulate organic matter (CPOM), scrapers feeding on autochthonous production (periphyton), collector-filterers feeding on suspended fine particulate organic matter (FPOM), collector-gatherers feeding on deposited FPOM and predators feeding on live animal tissue (Merritt et al., 2017). Shredders, scrapers and collectors were also subsumed as herbivores.

### 2.5 Data analysis

All physical and chemical data were transformed using natural-log transformations before analysis. We used one-way analysis of variance (ANOVA) to test for differences in water quality, stream size and land use variables among the five regions, followed by post hoc Tukey’s Honestly Significant Difference (HSD) multiple comparisons of means.

We used MixSIAR Bayesian mixing model to estimate the contributions of different basal sources to consumer diets (Parnell et al., 2010; Stock and Semmens, 2013). Models were run for each site separately for each FFG and common family. Only C3 and C4 plants and periphyton were included in the models as possible sources of carbon and nitrogen (Masese et al.; 2015, 2018). For periphyton, we calculated standard deviations of source values for input into MixSIAR from triplicate samples collected from the same site. Because the isotopic values of vegetation are influenced by elevation, for C3 and C4 sources we grouped sites into three groups defined by elevation: headwaters (> 2800 m a.s.l.), middle reaches (2000-2800 m a.s.l.) and lower reaches in the MMNR and surrounding areas (<2000 m a.s.l.). Trophic enrichment factors (TEFs) were set to zero after manually correcting for trophic fractionation in the stable isotope composition of consumers. For δ^15^N, we used a trophic fractionation of 0.6 ± 1.7 ‰ and 1.8 ± 1.7 ‰ for herbivorous and predatory macroinvertebrates, respectively (Bunn et al., 2013). For δ^13^C, we used a trophic fractionation of 0.5 ± 1.3 ‰ (McCutchan Jr et al., 2003). Concentration dependencies were set to zero.

For each site, river distance from the source (RDS) was calculated as the square root of the drainage area (Rasmussen et al., 2009) as a general measure reflecting the linear dimension of a watershed. This is based on the finding that the average length of stream paths leading to a point in the drainage can be expressed as a power function of the drainage area (Gregory and Walling, 1973) with the exponent 0.5 (Smart, 1972). We used RDS and LMH (livestock and hippos) density per sampling site as independent variables against which longitudinal changes in fractional contributions of periphyton, C3 and C4 carbon to invertebrates in the river were explored using simple linear regression (SLR). Relationships were tested separately for the different FFGs.

Data on livestock (cattle, sheep, goats, and donkeys) and wildlife (ungulates and hippos) numbers were obtained from Development Plans for Bomet and Narok Districts (Plan, 2007, 2008), Ministry of Agriculture and Livestock Production, and Kenya National Bureau of Statistics reports (KNBS-IHBS, 2007; KNBS-LS, 2009; KNBS, 2016, 2018), and other unpublished and published reports (Ottichilo et al., 2000; Lamprey and Reid, 2004; Kanga et al., 2011; Kiambi et al., 2012; Ogutu et al., 2016). For livestock and other LMH, density was expressed as the number of individuals per km^2^ in the catchment area of the river sampling site, while for hippos, count data were expressed as the number of individuals per river km.

We used generalized additive models (GAMs) (Wood, 2017) to assess how the relative importance of C3 carbon, C4 carbon and periphyton for individual FFGs and all FFGs combined varied with stream size (RDS) and density of LMH. GAMs incorporate smooth functions that are more flexible in modelling nonlinear relationships (Hastie and Tibshirani, 1990). Typically, the importance of different sources of carbon varies non-linearly with stream order or distance from the source (Vannote et al., 1980). GAMs were built using penalized cubic regression splines with degrees of freedom automatically identified based on the generalized cross-validation score (GCV) using the mgcv-package (Wood and Wood, 2015) in the R platform (R-Development-Core-Team, 2017). GAMs were used further to investigate the influence of region, stream size (RDS) and LMH density, including potential interactions, on the relative importance of the three sources of carbon for FFGs in the river. GAMs included region, RDS and LMH density as fixed effects, and individual sampling sites as a random effect. All analyses were performed using R 3.4.4 (Team, 2017).

## 3. Results

### 3.1 Physico-chemical variables

There was spatial variation in many physical and chemical variables (Table 1). LMH density increased with the proportion of forest cover and decreased with the proportion of grassland. Electrical conductivity and water temperature increased with LMH density but were low at the forested, low livestock density and agricultural regions/ sites (Table 1). Apart from the hippo sites, there were no differences in river distance from the source (RDS), river width, depth, or discharge among the five regions.

**Table 1.** River size, the density of large mammalian herbivores, catchment land use characteristics and δ^13^C and δ^15^N values of the main carbon sources (periphyton, C3 and C4 carbon, dung) across 80 sites of the Mara River, Kenya, grouped into five regions: Forested, Agricultural, with low density (LD) livestock, with high density (HD) livestock, and with a dominance of hippos. All values are means (± SD) and δ^13^C and δ^15^N values are in ‰ units.

| Characteristics | Regions |  |  |  |  | Statistics |  |
| --- | --- | --- | --- | --- | --- | --- | --- |
|  | Forested | Agricultural | LD livestock | HD livestock | Hippos | F - value | p - value |
| **Stream and sub-catchment characteristics** |  |  |  |  |  |  |  |
| Stream order | 4.0±1.5<sup>a</sup> | 3.7±1.6<sup>a</sup> | 4.8±2.1<sup>ab</sup> | 4.0±1.9<sup>a</sup> | 6.4±1.6<sup>b</sup> | 5.26 | 0.001 |
| River distance from source (RDS) | 12.5±6.4<sup>a</sup> | 12.2±9.2<sup>a</sup> | 26.0±17.4<sup>b</sup> | 17.2±15.8<sup>ab</sup> | 52.6±23.2<sup>b</sup> | 7.37 | 0.001 |
| LMH density (individuals / km<sup>2</sup>) | 9.3±4.1<sup>a</sup> | 28.0±16.8<sup>b</sup> | 44.8±12.5<sup>b</sup> | 102.9±17.7<sup>c</sup> | 111.9±10.8<sup>c</sup> | 11.21 | 0.001 |
| % Agriculture | 27.1±24.9<sup>a</sup> | 61.1±22.2<sup>b</sup> | 63.5±23.9<sup>b</sup> | 1.9±0.5<sup>c</sup> | 22.4±17.3<sup>ab</sup> | 18.91 | 0.001 |
| % Grasslands | 5.6±5.2<sup>a</sup> | 7.2±5.1<sup>a</sup> | 16.5±13.9<sup>b</sup> | 59.9±10.8<sup>c</sup> | 49.2±16.0<sup>c</sup> | 93.21 | 0.001 |
| % Forest | 64.7±28.5<sup>a</sup> | 39.3±22.9<sup>b</sup> | 25.4±15.4<sup>b</sup> | 37.2±11.5<sup>b</sup> | 27.97±8.9<sup>b</sup> | 9.47 | 0.001 |
| River width (m) | 7.9±5.0<sup>a</sup> | 7.1±6.1<sup>a</sup> | 8.7±5.8<sup>a</sup> | 6.6±3.5<sup>a</sup> | 15.84±9.9<sup>b</sup> | 4.35 | 0.003 |
| Water depth (m) | 0.20±0.1<sup>a</sup> | 0.2±0.1<sup>a</sup> | 0.2±0.2<sup>a</sup> | 0.2±0.1<sup>a</sup> | 1.0±0.4<sup>b</sup> | 5.45 | 0.001 |
| River discharge (m<sup>3</sup>/s) | 0.23±0.2<sup>a</sup> | 0.4±0.2<sup>a</sup> | 0.5±0.4<sup>a</sup> | 0.3±0.2<sup>a</sup> | 17.4±6.9<sup>b</sup> | 7.57 | 0.001 |
| pH (units) | 7.6±0.3<sup>a</sup> | 7.6±0.3<sup>a</sup> | 7.7±0.2<sup>a</sup> | 7.6±0.1<sup>a</sup> | 7.8±0.0<sup>a</sup> | 0.40 | 0.804 |
| Temperature (°C) | 15.8±1.8<sup>a</sup> | 18.5±3.2<sup>b</sup> | 20.3±2.5<sup>b</sup> | 23.7±2.1<sup>c</sup> | 23.9±2.2<sup>c</sup> | 27.9 | 0.001 |
| Dissolved oxygen (mgL<sup>-1</sup>) | 7.8±0.4<sup>a</sup> | 7.0±1.1<sup>bc</sup> | 6.3±0.9<sup>c</sup> | 7.3±1.0<sup>ab</sup> | 8.0±1.1<sup>a</sup> | 7.36 | 0.001 |
| Electrical conductivity ($\mu\text{Scm}^{-1}$) | 74.1±29.4<sup>a</sup> | 103.2±50.0<sup>a</sup> | 260.3±128.9<sup>b</sup> | 309.0±172.8<sup>b</sup> | 325.0±178.7<sup>b</sup> | 18.12 | 0.001 |
| **Stable isotopes of basal resources** |  |  |  |  |  |  |  |
| $\delta^{13}\text{C}$ of C3 vegetation | -26.56±2.08<sup>a</sup> | -27.14±3.37<sup>a</sup> | -27.62±1.37<sup>a</sup> | -25.36±1.98<sup>a</sup> | -26.13±1.97<sup>a</sup> | 1.05 | 0.407 |
| $\delta^{13}\text{C}$ of C4 vegetation | -13.22±6.27<sup>a</sup> | -12.08±1.56<sup>a</sup> | -16.44±7.95<sup>a</sup> | -11.04±2.13<sup>a</sup> | -15.64±11.01<sup>a</sup> | 1.32 | 0.294 |
| $\delta^{13}\text{C}$ of periphyton | -25.93±2.58<sup>a</sup> | -24.34±2.64<sup>ab</sup> | -23.7±2.15<sup>ab</sup> | -23.21±1.23<sup>b</sup> | -22.66±1.80<sup>b</sup> | 4.24 | 0.004 |
| $\delta^{13}\text{C}$ of cattle dung | -20.96±7.31<sup>a</sup> | -15.78±2.36<sup>ab</sup> | -13.49±1.73<sup>b</sup> | -14.27±5.01<sup>b</sup> | -13.58±2.85<sup>b</sup> | 5.08 | 0.002 |
| $\delta^{13}\text{C}$ of hippo dung | - | - | - | -11.89±2.14<sup>a</sup> | -14.13±2.72<sup>a</sup> | 2.11 | 0.169 |
| $\delta^{15}\text{N}$ of C3 vegetation | 4.06±1.69<sup>a</sup> | 5.47±4.96<sup>a</sup> | 5.24±2.65<sup>a</sup> | 4.61±2.36<sup>a</sup> | 3.37±2.91<sup>a</sup> | 1.83 | 0.119 |
| $\delta^{15}\text{N}$ of C4 vegetation | 6.09±2.59<sup>a</sup> | 5.98±3.07<sup>a</sup> | 4.76±4.79<sup>a</sup> | 5.57±3.25<sup>a</sup> | 4.81±2.74<sup>a</sup> | 0.15 | 0.988 |
| $\delta^{15}\text{N}$ of periphyton | 7.13±0.84<sup>a</sup> | 7.56±1.49<sup>ab</sup> | 8.53±1.44<sup>b</sup> | 8.11±0.79<sup>b</sup> | 7.29±0.71<sup>ab</sup> | 3.16 | 0.019 |
| $\delta^{15}\text{N}$ of cattle dung | 6.79±1.79<sup>a</sup> | 5.37±1.29<sup>a</sup> | 7.29±0.94<sup>a</sup> | 5.89±1.29<sup>a</sup> | 4.55±0.20<sup>a</sup> | 1.34 | 0.257 |
| $\delta^{15}\text{N}$ of hippo dung | - | - | - | 6.56±1.96<sup>a</sup> | 4.93±0.63<sup>a</sup> | 2.19 | 0.158 |

### 3.2 Basal resources

The C3 and C4 plants collected at the various sites did not differ significantly in δ^13^C and δ^15^N (Table 1), nevertheless mean isotopic signatures were calculated for each of three elevation levels for use in the MixSIAR models. The δ^15^N values of C4 plants were generally higher than those of C3 plants. δ^13^C, but not δ^15^N, of cattle dung, varied considerably among sites and regions (Table 2). In contrast, δ^13^C and δ^15^N of hippo dung showed little variation among regions with hippos. δ^13^C and δ^15^N of periphyton were highly variable among sites and regions (Figure 2, Table 1); specifically, agricultural and livestock sites had elevated δ^15^N values compared to forested areas and the MMNR, where hippos had a strong influence (Figure 2). In MixSIAR models, site-specific isotopic signatures were used for all resources with detectable variation among regions.

**Table 2.**
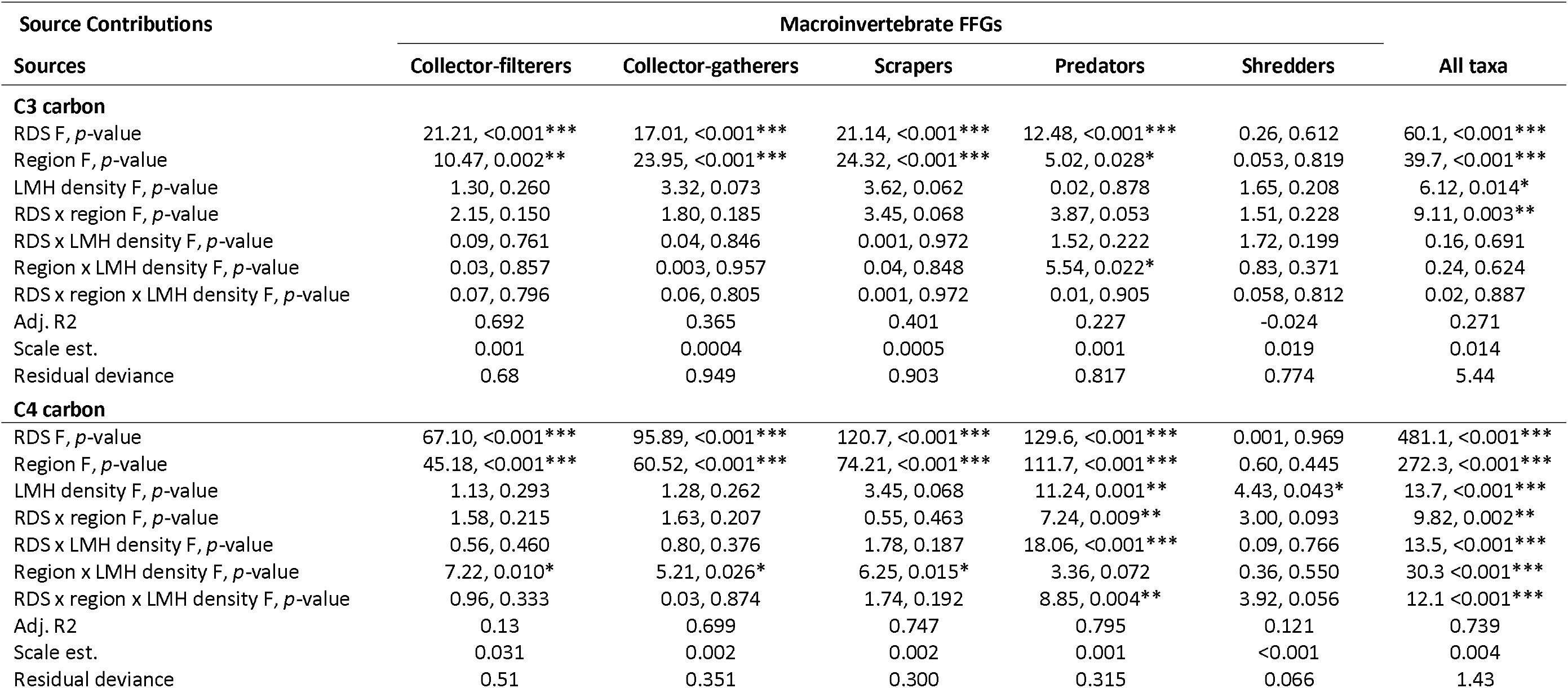
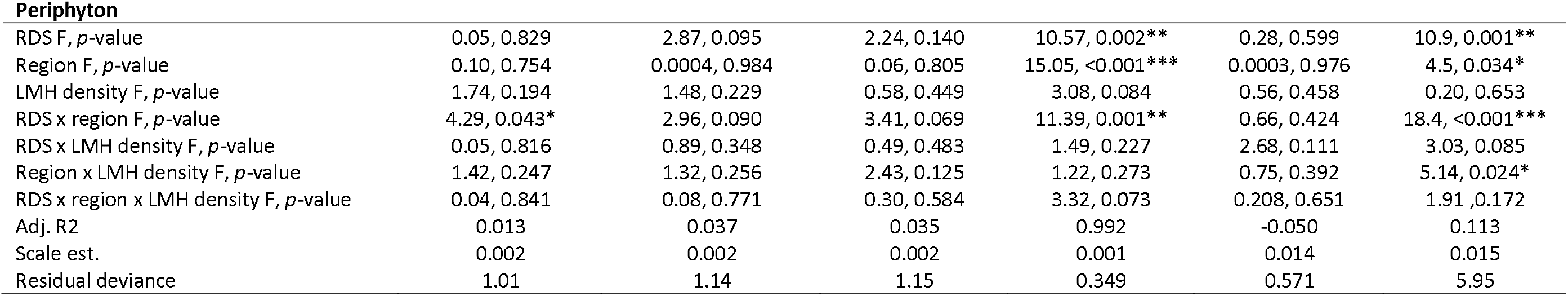
Summary of generalized additive mixed models (GAMs) of the effects of stream size (river distance from source {RDS}), region and density of large mammalian herbivores (LMH density) (stream size) on the relative importance (fractional contribution) of C3 vegetation (C3 carbon), C4 grasses (C4 carbon) and periphyton as sources of carbon/energy for macroinvertebrate FFGs (collector-filterers, collector-gatherers, scrapers, predators and shredders) and all combined taxa (FFGs) in the Mara River, Kenya. RDS, region and LMH density were treated as fixed factors while each study site was used a random factor. In all cases, d.f. = 1.

| Source Contributions |  | Macroinvertebrate FFGs |  |  |  |  |
| --- | --- | --- | --- | --- | --- | --- |
| Sources | Collector-filterers | Collector-gatherers | Scrapers | Predators | Shredders | All taxa |
| **C3 carbon** |  |  |  |  |  |  |
| RDS F, *p*-value | 21.21, <0.001*** | 17.01, <0.001*** | 21.14, <0.001*** | 12.48, <0.001*** | 0.26, 0.612 | 60.1, <0.001*** |
| Region F, *p*-value | 10.47, 0.002** | 23.95, <0.001*** | 24.32, <0.001*** | 5.02, 0.028* | 0.053, 0.819 | 39.7, <0.001*** |
| LMH density F, *p*-value | 1.30, 0.260 | 3.32, 0.073 | 3.62, 0.062 | 0.02, 0.878 | 1.65, 0.208 | 6.12, 0.014* |
| RDS x region F, *p*-value | 2.15, 0.150 | 1.80, 0.185 | 3.45, 0.068 | 3.87, 0.053 | 1.51, 0.228 | 9.11, 0.003** |
| RDS x LMH density F, *p*-value | 0.09, 0.761 | 0.04, 0.846 | 0.001, 0.972 | 1.52, 0.222 | 1.72, 0.199 | 0.16, 0.691 |
| Region x LMH density F, *p*-value | 0.03, 0.857 | 0.003, 0.957 | 0.04, 0.848 | 5.54, 0.022* | 0.83, 0.371 | 0.24, 0.624 |
| RDS x region x LMH density F, *p*-value | 0.07, 0.796 | 0.06, 0.805 | 0.001, 0.972 | 0.01, 0.905 | 0.058, 0.812 | 0.02, 0.887 |
| Adj. R2 | 0.692 | 0.365 | 0.401 | 0.227 | -0.024 | 0.271 |
| Scale est. | 0.001 | 0.0004 | 0.0005 | 0.001 | 0.019 | 0.014 |
| Residual deviance | 0.68 | 0.949 | 0.903 | 0.817 | 0.774 | 5.44 |
| **C4 carbon** |  |  |  |  |  |  |
| RDS F, *p*-value | 67.10, <0.001*** | 95.89, <0.001*** | 120.7, <0.001*** | 129.6, <0.001*** | 0.001, 0.969 | 481.1, <0.001*** |
| Region F, *p*-value | 45.18, <0.001*** | 60.52, <0.001*** | 74.21, <0.001*** | 111.7, <0.001*** | 0.60, 0.445 | 272.3, <0.001*** |
| LMH density F, *p*-value | 1.13, 0.293 | 1.28, 0.262 | 3.45, 0.068 | 11.24, 0.001** | 4.43, 0.043* | 13.7, <0.001*** |
| RDS x region F, *p*-value | 1.58, 0.215 | 1.63, 0.207 | 0.55, 0.463 | 7.24, 0.009** | 3.00, 0.093 | 9.82, 0.002** |
| RDS x LMH density F, *p*-value | 0.56, 0.460 | 0.80, 0.376 | 1.78, 0.187 | 18.06, <0.001*** | 0.09, 0.766 | 13.5, <0.001*** |
| Region x LMH density F, *p*-value | 7.22, 0.010* | 5.21, 0.026* | 6.25, 0.015* | 3.36, 0.072 | 0.36, 0.550 | 30.3 <0.001*** |
| RDS x region x LMH density F, *p*-value | 0.96, 0.333 | 0.03, 0.874 | 1.74, 0.192 | 8.85, 0.004** | 3.92, 0.056 | 12.1 <0.001*** |
| Adj. R2 | 0.13 | 0.699 | 0.747 | 0.795 | 0.121 | 0.739 |
| Scale est. | 0.031 | 0.002 | 0.002 | 0.001 | <0.001 | 0.004 |
| Residual deviance | 0.51 | 0.351 | 0.300 | 0.315 | 0.066 | 1.43 |
| **Periphyton** |  |  |  |  |  |  |
| RDS F, *p*-value | 0.05, 0.829 | 2.87, 0.095 | 2.24, 0.140 | 10.57, 0.002** | 0.28, 0.599 | 10.9, 0.001** |
| Region F, *p*-value | 0.10, 0.754 | 0.0004, 0.984 | 0.06, 0.805 | 15.05, <0.001*** | 0.0003, 0.976 | 4.5, 0.034* |
| LMH density F, *p*-value | 1.74, 0.194 | 1.48, 0.229 | 0.58, 0.449 | 3.08, 0.084 | 0.56, 0.458 | 0.20, 0.653 |
| RDS x region F, *p*-value | 4.29, 0.043* | 2.96, 0.090 | 3.41, 0.069 | 11.39, 0.001** | 0.66, 0.424 | 18.4, <0.001*** |
| RDS x LMH density F, *p*-value | 0.05, 0.816 | 0.89, 0.348 | 0.49, 0.483 | 1.49, 0.227 | 2.68, 0.111 | 3.03, 0.085 |
| Region x LMH density F, *p*-value | 1.42, 0.247 | 1.32, 0.256 | 2.43, 0.125 | 1.22, 0.273 | 0.75, 0.392 | 5.14, 0.024* |
| RDS x region x LMH density F, *p*-value | 0.04, 0.841 | 0.08, 0.771 | 0.30, 0.584 | 3.32, 0.073 | 0.208, 0.651 | 1.91, 0.172 |
| Adj. R2 | 0.013 | 0.037 | 0.035 | 0.992 | -0.050 | 0.113 |
| Scale est. | 0.002 | 0.002 | 0.002 | 0.001 | 0.014 | 0.015 |
| Residual deviance | 1.01 | 1.14 | 1.15 | 0.349 | 0.571 | 5.95 |

**Figure 2.**
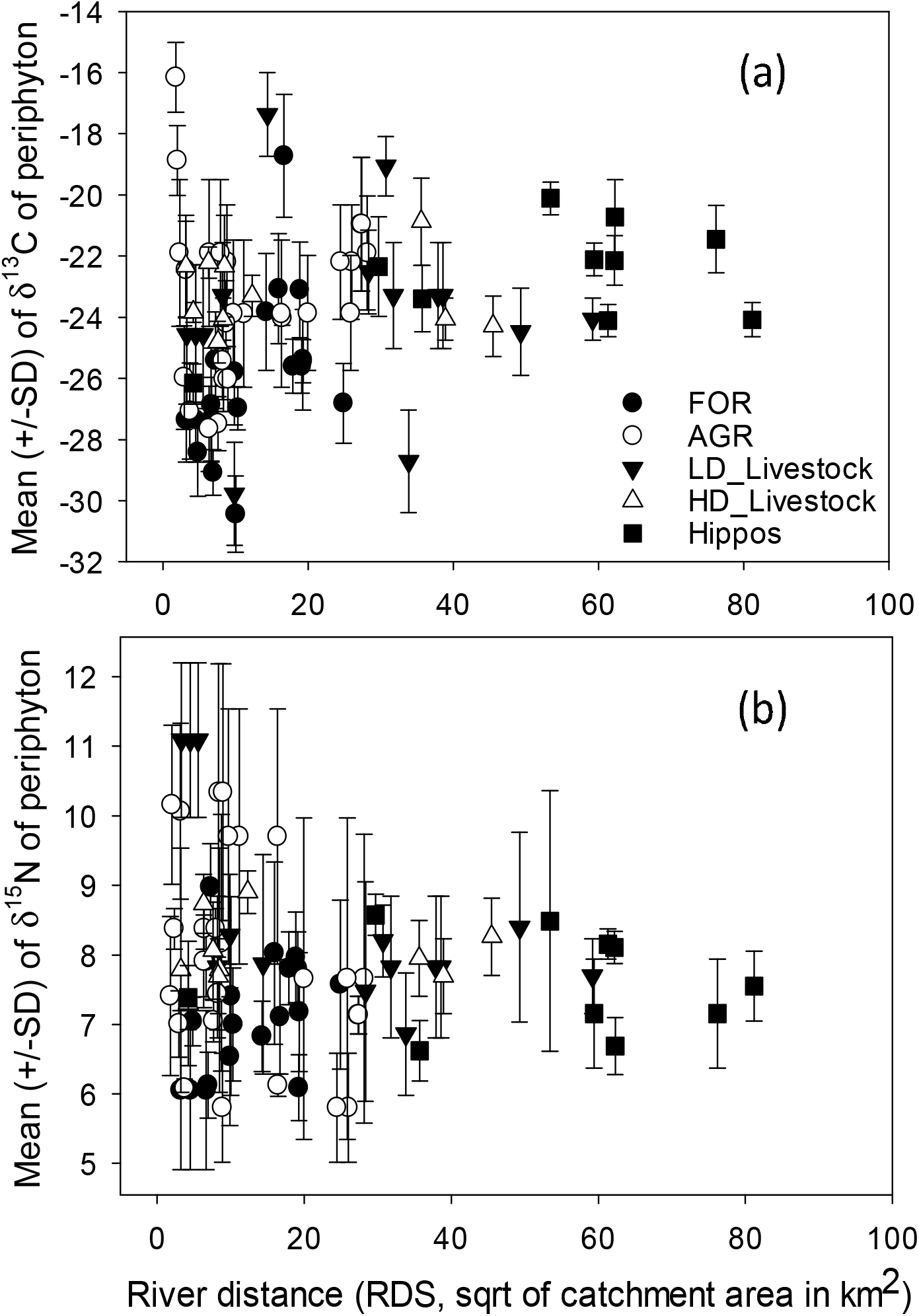
Longitudinal trends in the mean (±SD) of (a) δ^13^C and (b) δ^15^N values (‰) of periphyton with increasing stream size in the Mara River and its tributaries. FOR = forested sites, AGR = agriculture sites, LD_Livestock = low density livestock sites, HD_Livestock = high density livestock sites, and Hippos = hippopotamus sites.

### 3.3 Longitudinal trends in basal resources

Longitudinal patterns in δ^13^C of periphyton in the Mara River were wedge-shaped (Figure 2). Periphyton δ^13^C responded to both measures of stream size RDS (r = 0.23, p < 0.01) and stream order (r = 0.35, p < 0.01). At shorter RDS, agricultural and low density (LD) livestock sites had the widest range of periphyton δ^13^C, while values at forested sites were invariably low (−25.9±2.6‰, Table 2). The narrowest ranges of δ^13^C were recorded for high density (HD) livestock and hippo sites, which appeared at longer RDS. At these sites, δ^13^C (HD livestock: −23.2±1.2‰, hippo sites: −22.7±1.8‰) was on average higher than at forested sites. δ^13^C of periphyton was positively related to agricultural land use (r = 0.24, p < 0.05) and negatively to forest cover (r = −0.44, p < 0.01). Overall, periphyton δ^13^C responded positively to LMH density (r = 0.52, p < 0.01), which itself was highly positively correlated with % grassland cover in the catchment (r = 0.89, p < 0.01).

Longitudinal patterns of periphyton δ^15^N were similar to δ^13^C, yet we could not detect a significant change of the mean value with any measure of stream size. Periphyton δ^15^N was higher at agricultural and LD livestock sites (Figure 2) with shorter RDS, but lower at both the forested sites with short RDS and hippo sites at longer RDS. Unexpectedly, δ^15^N at LD livestock sites also surpassed HD livestock sites. δ^15^N of periphyton was positively related to agricultural land use (r = 0.29, p < 0.01) and negatively to forest cover (r = −0.31, p < 0.01). Interestingly, there was no significant relationship between δ^15^N of periphyton and LMH density.

### 3.4 Importance of different carbon sources for invertebrates

A total of 47 taxa in the five macroinvertebrate FFGs were collected in the study area (Table S1). The importance of C3 vegetation, C4 grasses and periphyton for invertebrates differed among the five regions, but patterns were similar for scrapers and collectors (Figure 3). Overall, periphyton was either the major or second-most important source of carbon for all FFGs at forested, agricultural and livestock sites, and predators were more reliant on this energy pathway than the rest of the FFGs (Figure 3). Except for shredders in forested streams, the importance of C3 vegetation was reduced for the rest of FFGs in the other regions and was lowest at the hippo sites. On the contrary, the importance of C4 carbon responded strongly to LMH density and was the most important source of carbon (>50%) for all FFGs at hippo sites, except shredders (Figure 3).

**Figure 3.**
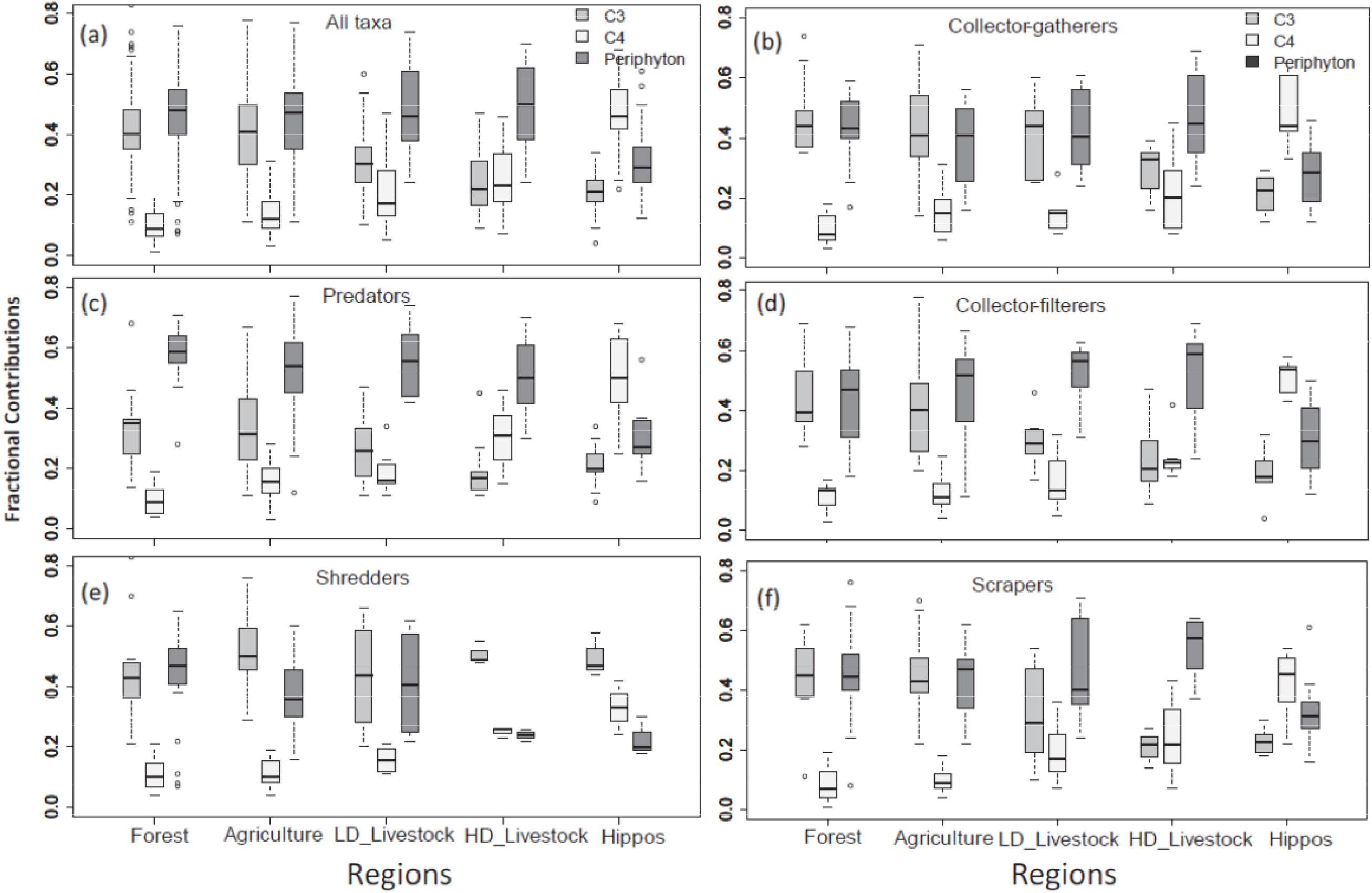
Box-and-whisker plots of fractional contributions of C3 vegetation (grey), C4 grasses (light grey) and periphyton (dark grey) to all macroinvertebrates FFGs in the Mara River. (a) all taxa, (b) collector-gatherers, (c) predators, (d) collector-filterers (e) shredders, and (f) scrapers. C3 = C3 plants; C4 = C4 plants.

Source contributions followed similar patterns for all FFGs except predators and shredders (Figure 3). Predators seemed to draw most of their nutrition from the periphyton pathway at all sites except those with hippos. Shredders, on the other hand, displayed an overreliance on C3 vegetation as their main source of carbon at all sites, which is consistent with their feeding habits on coarse particulate organic matter, i.e., mostly leaves of C3 vegetation of terrestrial origin. Thus, shredders were least influenced by inputs of C4 carbon into the river via LMH.

### 3.5 Longitudinal trends in the importance of different carbon sources

We explored relationships between stream size (i.e., river distance from the source, RDS) and LMH density as predictors of the importance of the three differentiated sources of carbon (C3 vegetation, C4 grasses and periphyton) for macroinvertebrates in the Mara River (Figure 4). The relative importance (fractional contribution) of the three sources of carbon changed with stream size (RDS) and LMH density (Figure 4a, b). Periphyton was the most important source of energy in headwater and mid-sized streams, contributing on average more than 40% to invertebrate biomass, but decreased to the second most important source of carbon further downstream. C3 vegetation (woody vegetation) was almost as important as periphyton in headwater streams, contributing nearly 40%, but its importance declined from mid-sized streams onwards, becoming the least important source of carbon with contributions <20% in large river sections (Figure 4a). C4 plants (grasses) were not relevant in small streams contributing less than 20% to invertebrates’ energy requirements, but their importance markedly increased in mid-sized and large river sections, where they became the dominant source of energy with a contribution of nearly 60% to the total energy requirements for macroinvertebrates. The influence of LMH on the longitudinal importance of carbon sources for macroinvertebrates in the river (Figure 4b) mirrored that of RDS, although patterns for C4 carbon were stronger. Patterns with stream order as a proxy for stream size were identical to those with RDS (data not shown).

**Figure 4.**
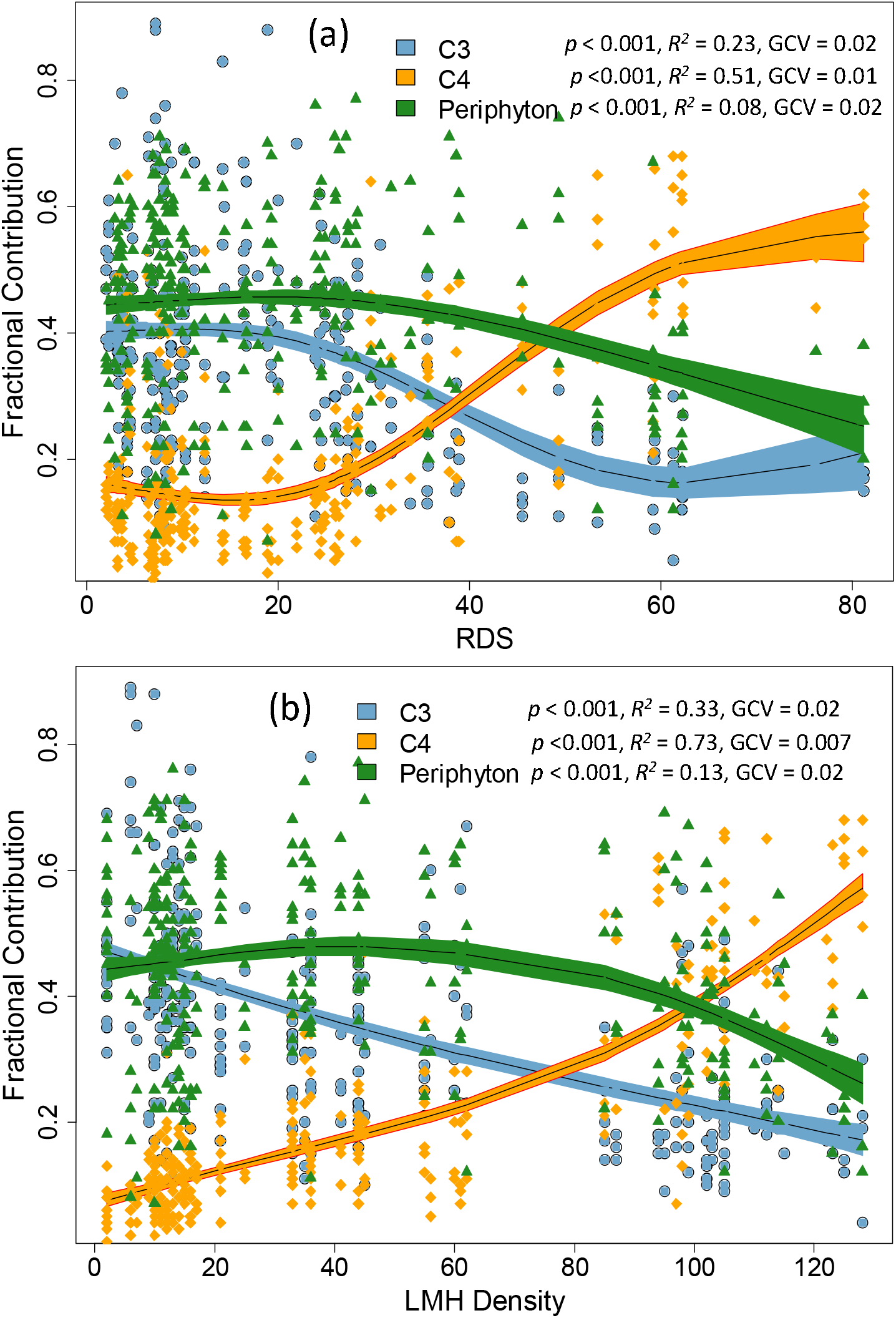
Longitudinal trends in source contributions to macroinvertebrates along the Mara River. Longitudinal variability in the relative importance (fractional contribution) of C3 vegetation (sky blue), C4 grasses (orange) and periphyton (dark green) as sources of carbon for macroinvertebrates in the Mara River in response to changes in (a) river distance from the source (RDS in km) and (b) density of large mammalian herbivores (LMH Density). To test the significance of the relationships, we fitted a GAM model with a smoothing function. The black line with shaded area represents smoother mean and s.e.; smoother significance, *R*^2^ and GCV are supplied in the figures.

We also explored longitudinal relationships between stream size (i.e., river distance from the source, RDS) and LMH density as predictors of the importance of the three sources of carbon (C3 vegetation, C4 grasses and periphyton) for individual FFGs (Figure 5). There were significant positive relationships (linear regressions, p < 0.05) between RDS and the importance of C4 to all FFGs except shredders (Figure 5g, h). In parallel, C3 carbon from terrestrial vegetation became less important with RDS for collector-filters (Figure 5a, b) and collector-gatherers (Figure 5c, d). Also, LMH density had strong influences on the importance of carbon sources for all FFGs except shredders, by reducing the contribution of C3 while increasing that of C4 carbon. As expected, shredders were mainly reliant on C3 vegetation and were neither influenced by stream size nor LMH density (Figure 5g, h).

**Figure 5.**
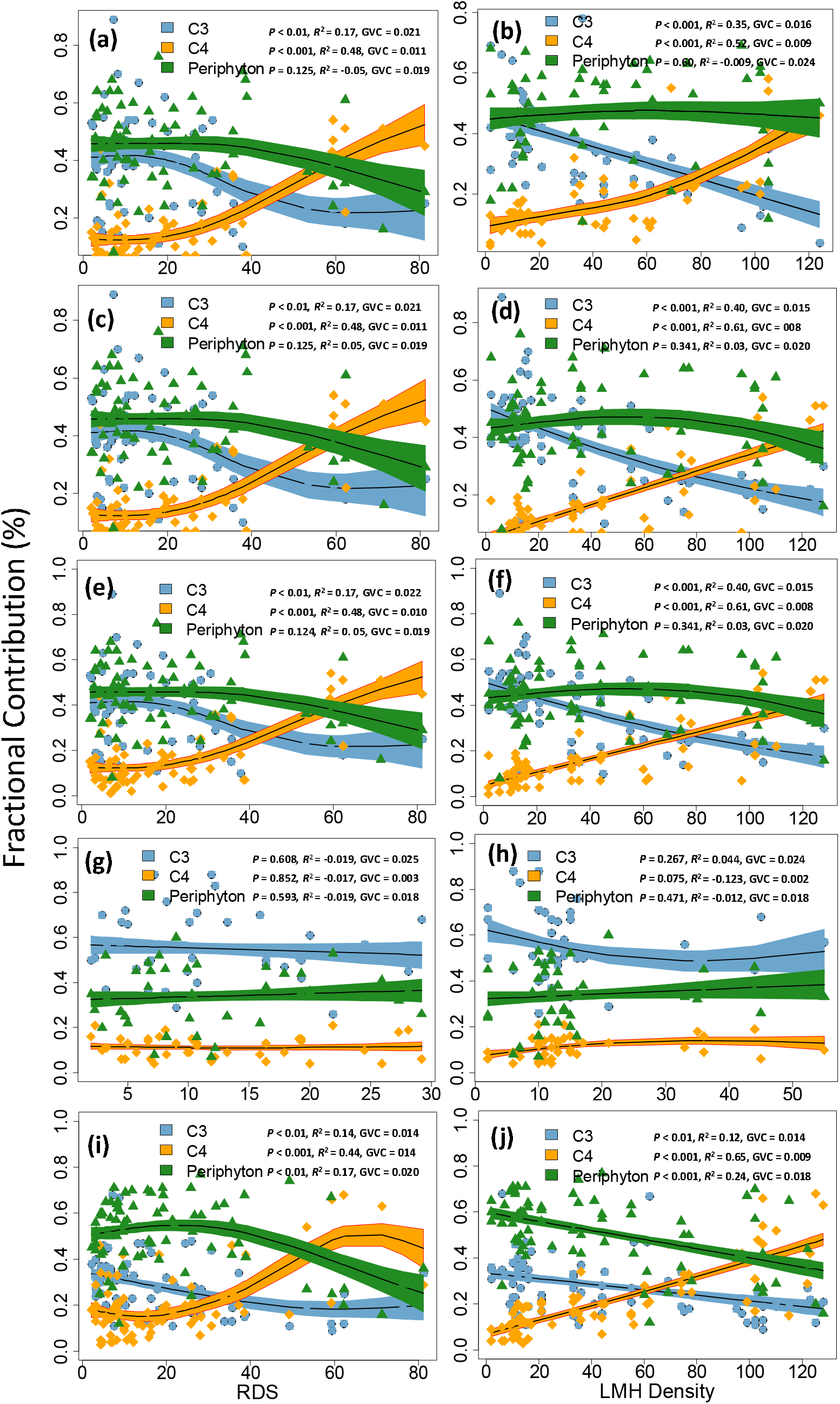
Source contributions to FFGs along the Mara River. Longitudinal trends in fractional contributions of trophic source (C3= C3 vegetation, C4 = C4 grasses and periphyton) to macroinvertebrate functional feeding groups (FFGs) in the Mara River. Source contributions are assessed in response to changes in river distance from source (RDS) as a measure of stream size (a, c, e, g, i) and density of large mammalian herbivores (LMH), b, d, f, h, j). a, b = collector-filterers, c, d = collector-gatherers, e, f = scrapers, g, h = shredders, and I, j = predators. To test the significance of the relationships, we fitted a GAM model with a smoothing function. The black line with shaded area represents smoother mean and s.e.; smoother significance, *R*^2^ and GCV are supplied in the figures. Note changes on the x-axis and y-axis.

We found a significant influence of stream size and region on the relative importance of C3 and C4 carbon for combined and individual FFGs, except shredders, in the river (Table 2). The independent effects of LMH density on the relative importance of C4 carbon was limited to scrapers and predators only, implying increasing LMH density did not necessarily increase the importance of C4 carbon for invertebrates. This is in agreement with observations that livestock increased the importance of periphyton while hippos increased the importance of C4 carbon for most FFGs, but increasing densities of both livestock and hippos decreased the importance of C3 carbon for invertebrates (Figure 3). There were significant interactions between LMH density and region in the importance of C4 carbon for all FFGs, except shredders. Overall, patterns in the importance of the three sources of carbon for predators were similar to those of all FFGs combined.

To eliminate potential confounding by organic matter and nutrient inputs by other agricultural activities not associated with LMH, the influence of stream size and LMH density on the relative importance of carbon sources for invertebrate FFGs were evaluated in forested sites only where LMH (both livestock and wildlife) were limited (Supplementary Information, Figures S1). Remarkably, no major effect of stream size was noted for most FFGs, further reinforcing the important role of LMH as drivers of food resources for macroinvertebrates in the river.

## 4. Discussion

This study shows that the importance of different sources of carbon for consumers in savanna rivers is spatially variable in response to changes in abundance and distribution of large mammalian herbivores, both livestock and wildlife, and stream size or river distance from the source. Overall, (i) periphyton was the most important source of energy for macroinvertebrates in low order and mid-sized rivers, (ii) periphyton and C4 vegetation became more important carbon sources at the cost of C3 plants at high-density livestock and hippo sites, respectively, and (iii) lower sections of the river hosting large populations of LMH relied largely on C4 vegetation with very little contributions of C3 plants. Our findings thus confirm our hypothesis of an overriding influence of LMH over stream size or location along the fluvial continuum in determining sources of carbon for food webs in savanna rivers.

Although previous studies in African savanna rivers have shown that LMH foster the importance of C4 carbon as a major source of carbon for aquatic consumers (Masese et al. 2015, McCauley et al., 2015, Dawson et al., 2020), no study has evaluated the importance of these resource subsidies on entire gradients of rivers and at the scale that we have done in this study. The many stream sites sampled that differed in size and LMH density enabled us to evaluate the interaction of stream size with subsidy quantity as determinants for the base of riverine food webs. Our results depart from predictions on the importance of resources along the river continuum. While a reduction in the contribution of C3 carbon is hypothesized in mid-sized rivers as periphyton becomes dominant (Vannote et al., 1980; Thorp and Delong, 1994), an increase in the importance of fresh C4 vegetation inputs to mid-sized and large rivers is less intuitive and not hypothesized in existing models of riverine ecosystem functioning. Instead, mid-sized and large rivers are thought to be disproportionately reliant on autochthonous production, and leakage of unutilized materials from upstream supply food webs of large rivers (Vannote et al., 1980, Thorp and Delong, 2002; Thorp et al., 2006).

Although LMH density and the proportion of C4 grasses in the catchments were strongly correlated (r = 0.89), the increased importance of C4 carbon in the diet of most macroinvertebrates in this study was assumed to be mediated via the vectoring role of LMH when they defecate in the river. Indeed, in the absence of LMH, the contribution of C4 carbon to riverine food webs is less than expected based on areal landcover (Abrantes and Sheaves, 2010; Marwick et al., 2014), and given lower C4 vegetation height direct riparian inputs were minimal. Without LMH-mediation, C4 carbon would hardly reach aquatic food webs and dominance of C3 carbon could be expected in larger rivers, either supplied via leakage from upstream or through direct deposition from riparian vegetation. Headwater streams and riparian areas of most savanna rivers contain more C3 than C4 vegetation (Table 1), implying that LMH are the critical vectors for C4 carbon, especially during the dry season.

Although both livestock and hippos largely facilitated the disproportionate contribution of C4 carbon from grasses to invertebrates in the river, there were interesting differences in the contribution of other sources (C3 vegetation and periphyton) between sites receiving inputs by livestock and hippos, which can be linked to differences in the quality, amount and characteristics of the inputs these two types of LMH transfer into rivers (Iteba et al., 2021; Masese et al., 2020). While increasing livestock density increased both the importance of periphyton and C4 carbon for macroinvertebrates in the river, hippo populations only increased the importance of C4 carbon further and reduced that of periphyton further (Figure 3). Although hippo dung has been reported to fertilize aquatic ecosystems increasing primary and secondary production (Grey and Harper, 2002; Mosepele et al., 2009), other studies have shown that large particles in hippo dung can have detrimental effects on benthic production, especially during the dry season when they settle at the bottom of hippo pools and downstream sections of rivers (Dawson et al., 2016; Dutton et al., 2018a, b). Hippos also spend long periods in the water for thermoregulation, approximately 12 h of daytime (Subalusky et al., 2015), and their wallowing activity is known to increase turbidity that can further limit primary production (Dutton et al., 2018). Comparative studies with cattle dung have also shown that the weakly digested hippo dung with high C: nutrients (N and P) ratios support heterotrophic microbial activity, while cattle dung with lower stoichiometric ratios can increase primary production faster (Subalusky et al., 2018; Masese et al., 2020). Thus, replacing wildlife (hippopotamus) with livestock (cattle) in savanna landscapes will most likely weaken detrital food web links based on allochthonous C3 and C4 material (Dawson et al., 2020) and foster an algal-grazer food chain by increasing the abundance of autochthonous material.

The proportional acquisition of energy from terrestrial C3 and C4 plants and periphyton showed specific patterns for different FFGs (Zeug and Winemiller, 2008; Pingram et al., 2014). Further, the importance of different sources was influenced by land use and the density of LMH. For instance, in forested streams, there was no effect of stream size on the relative importance of the three sources (C3 and C4 producers and periphyton) for most of the FFGs (Figure S1). There was significant partitioning among FFGs on the utilization of various resources, with scrapers and predators relying more on periphyton while collector-gatherers and, as expected, shredders relying more on C3 vegetation (Figure 1). However, facultative shredders, such as Potamonautidae (freshwater crabs) that dominated invertebrates most forested and agricultural streams, can obtain a significant amount of energy from periphyton during their omnivorous feeding (Masese et al., 2014). The importance of trophic sources was also related to the trophic position of the macroinvertebrates, with autochthonous production becoming more important with an increase in trophic level (Figures S1).

### 4.1 Longitudinal trends of the relative importance of trophic resources

Even in the absence of LMH (both livestock and wildlife) longitudinal patterns of organic matter and nutrient loading into the Mara River and its tributaries (Figures 4 and 5) departed significantly from existing predictions for river continuum (Vannote et al., 1980; Thorp and Delong, 2002; Thorp et al., 2006). Periphyton, and not leaf litter from C3 vegetation, was the most important source of energy for invertebrates in low-order forested streams by contributing nearly 50%, on average, of the energy requirements of all invertebrates while C3 vegetation contributed no more than 40%. These results have been supported by previous studies in tropical and temperate regions whereby carbon from autochthonous production contributes significantly to aquatic animal biomass in mid-sized and large rivers despite forming a small proportion of available food resources (Douglas et al., 2005; Hayden et al., 2016).

The importance of C3 vegetation for invertebrates declined predictably in mid-sized rivers, but an expected rebound in lower reaches of the river where turbidity limits primary production and the river becomes heterotrophic with an overreliance on fine particulate organic matter, mainly from C3 vegetation, escaping from upstream (Vannote et al., 1980), did not occur. Instead, the importance of C3 carbon further plummeted, while that of C4 carbon, mediated by large populations of livestock and large wildlife (mainly hippos), gained more importance. Moreover, the expected increase in the importance of autochthonously produced carbon in large rivers (Thorp and Delong, 1994, 2002) did not occur, further highlighting the predominant role played by LMH as drivers of ecosystem productivity and functioning in savanna rivers (Stears et al., 2018; Subalusky et al., 2018; Dawson et al., 2020; Masese et al., 2020).

Recognizing that carbon flow in riverine food webs is context-dependent, both temporally in terms of flow variability and spatially in terms of the strength of lateral and longitudinal connectivity, it is conceivable that the patterns we report here are transient. However, studies during both the dry and wet seasons, have indicated the overriding influence of LMH inputs on river water quality and biogeochemistry (Stears et al., 2018; Dutton et al., 2018b, 2020), ecosystem productivity (Subalusky et al., 2017, 2018), community composition and diversity of invertebrates and fishes (Masese et al., 2018; Stears et al., 2018). The discrepancies between the results of this study, on the relative importance of different sources of carbon for supporting riverine food webs on the fluvial continuum of savanna rivers, and existing models of riverine ecosystem functioning suggests that a new model is needed that recognizes the intimate linkage between terrestrial and aquatic ecosystems in savanna landscapes mediated by the active transfer of organic matter by LMH.

## 5. Conclusions

An understanding of terrestrial-aquatic ecosystems connectivity and the main sources of energy supporting riverine food webs is critical for the conservation and management of African savanna rivers. The novel findings of this study show the distinct functioning of these rivers in terms of the major energy sources fuelling food webs along their riverine continuum and highlight the important role played by LMH as vectors enhancing aquatic productivity through strengthened terrestrial-aquatic subsidy fluxes. We show that terrestrial and autochthonous sources of energy are relatively important along the longitudinal gradient of savanna rivers, and identify LMH as critically enhancing the contribution of C4 carbon far above potential contributions in their absence. The findings of this study also show that different resources are partitioned among macroinvertebrates depending on their feeding modes and location on the fluvial continuum. At a local scale, this study presents further evidence on the species-specific influences of different species of LMH on the functioning of aquatic ecosystems in African savannas. It also suggests that replacing native populations of large herbivores (such as hippos) with livestock (such as cattle) may alter the major sources of energy supporting aquatic communities in savanna rivers, with consequences on ecosystem structure and functioning.

## Acknowledgements

We are grateful to Mary Kiplagat (University of Eldoret) and Evans Ole Keshe (Mario Tours) for assistance in the field. We are grateful to Tobias Goldhammer, Sarah Krocker and Claudia Schmalsch for help during the analysis of water chemistry samples at IGB, Berlin. We thank Susanne Remus for stable isotope analysis at the ZALF. We thank Narok County for granting us access to the Mara Triangle. This is a publication of the MaMeMa Project and was funded, in part, by an Alexander von Humboldt Postdoc fellowship to FOM; and an ERC Research grant to GS.

## Notes

### Competing Interest Statement

The authors have declared no competing interest.

