## Supplementary Table S1 and Figure S1 for "Large herbivorous wildlife and livestock differentially influence the relative importance of different sources of carbon for riverine food webs"

***for***

**Macroinvertebrate FFGs analysed**

A total of 47 taxa belonging to different macroinvertebrate FFGs were collected in the study area (Table S1). For small-bodied taxa, different numbers of individuals were used to obtain enough material used for stable isotope analysis, while single individuals were used for the large-bodied invertebrates, such as freshwater crabs and most odonates.

Table S1. The number of invertebrate samples (n) analysed for δ^13^C and δ^15^N values or each taxon and functional feeding groups (FFGs) across 80 sites in the Mara River, Kenya, grouped into 5 categories: Forested, Agricultural, low density (LD) livestock, high density (HD) livestock and hippos.

|  |  | **Site categories** | | | | |
| --- | --- | --- | --- | --- | --- | --- |
| **FFGs** | **Taxa** | **Forested** | **Agriculture** | **LD Livestock** | **HD Livestock** | **Hippos** |
| Filterers | Bivalvia |  | 4 | 2 |  | 1 |
|  | Hydropsychidae | 29 | 42 | 26 | 15 | 9 |
|  | Simuliidae | 3 | 16 | 1 | 11 | 4 |
| Gatherers | Baetidae | 26 | 32 | 14 | 9 | 13 |
|  | Caenidae |  | 2 | 2 | 7 | 4 |
|  | Chironomidae | 3 | 6 | 3 | 7 | 6 |
|  | Dixidae |  |  |  |  | 2 |
|  | Lumbriculidae | 2 | 2 |  |  |  |
|  | Muscidae |  | 1 |  |  |  |
|  | Syriphidae |  |  |  |  | 1 |
| Predators | Aeshnidae | 6 | 5 |  | 2 | 1 |
|  | Arachnida |  |  |  | 2 |  |
|  | Belostomatidae |  | 3 | 1 |  |  |
|  | *Centroptiloides* sp. | 18 | 6 |  |  | 2 |
|  | Ceratopogonidae |  | 1 | 1 |  |  |
|  | Cordulliidae |  |  |  | 3 |  |
|  | Dytiscidae | 2 |  | 1 | 3 | 4 |
|  | Gerridae |  | 4 | 2 |  |  |
|  | Gomphidae | 38 | 26 | 12 | 8 | 11 |
|  | Gyrinidae |  | 4 | 3 |  | 4 |
|  | Helophoridae |  |  |  |  | 2 |
|  | Lestidae | 28 | 49 | 26 | 13 | 12 |
|  | Libellulidae |  | 2 | 2 |  |  |
|  | Naucoridae | 1 | 14 | 13 | 10 | 9 |
|  | Nepidae | 4 | 5 | 8 | 6 |  |
|  | Noteridae |  | 1 |  | 5 | 6 |
|  | Notonoctidae | 4 | 2 | 3 | 5 |  |
|  | Perlidae | 20 | 4 | 6 |  | 1 |
|  | Tabanidae |  |  |  | 1 | 3 |
| Scrapers | Heptageniidae | 42 | 51 | 19 | 5 | 8 |
|  | Baetidae 2 | 5 | 8 | 4 |  |  |
|  | Caenidae |  |  |  |  | 2 |
|  | Corixidae |  |  | 1 | 6 | 3 |
|  | Elmidae | 1 | 1 | 1 | 2 | 6 |
|  | Leptophlebiidae | 3 |  |  |  |  |
|  | Gastropoda |  |  | 1 |  |  |
|  | Philopotamidae | 3 |  |  |  |  |
|  | Tricorythidae | 4 | 5 | 2 | 2 | 1 |
| Shredders | Baetidae sp. | 3 | 3 |  |  |  |
|  | Crambidae | 16 | 13 | 2 | 1 |  |
|  | Larainae | 5 |  | 4 |  |  |
|  | Lepidostomatidae | 16 | 28 | 1 |  |  |
|  | Leptoceridae | 13 |  |  |  |  |
|  | Potamonautidae | 30 | 19 | 2 |  |  |
|  | Tipulidae | 3 | 13 | 2 | 1 |  |

**Potential for confounded effects of LMH density and human activities on importance of sources of carbon for food webs along the river**

To eliminate confoundment by large mammalian herbivore (LMH) density and stream size (RDS) on the relative importance of C3 and C4 carbon and periphyton to macroinvertebrates along the Mara River, relationships were evaluated for forested sites only where the number of LMH was low (Figure S1). Remarkably, limited to no effect of RDS was noted on the relative importance of the three sources of carbon for all FFGs, further reinforcing the role of LMH as drivers of the importance of food resources for macroinvertebrates along the river. Similarly, there was no influence of stream size on the relative importance of C3 vegetation, C4 grasses and periphyton to all FFGs in forested streams (Figures S1).

For all FFGs in forested streams, C4 carbon was the least important resource, while the importance of C3 vegetation and periphyton depended on the FFG in consideration. For instance, periphyton and C3 were equally important of scrapers (Figure S1e, f), C3 vegetation (woody vegetation) was the main source of carbon for shredders (Figure S1g, h). Interestingly, periphyton was the major source of carbon for predators (Figure S1i, j).

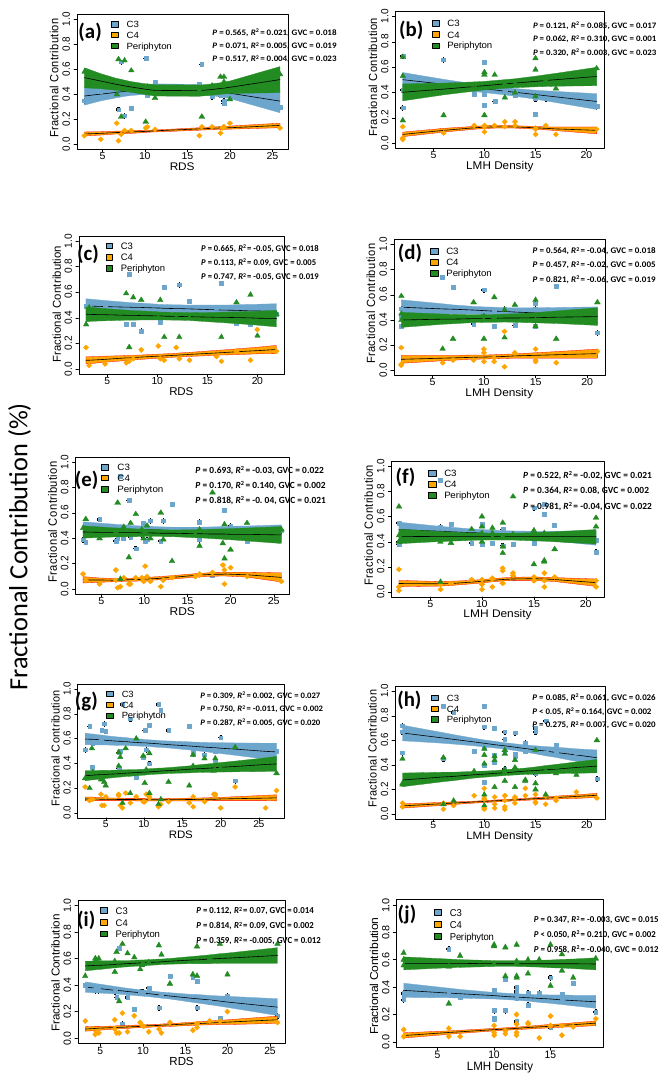

**Figure S1. Source contributions to FFGs in forested sites only.** Longitudinal trends in the fractional contribution of the three sources of carbon/ energy (C3 vegetation, C4 grasses and periphyton/ algae) to macroinvertebrate functional feeding groups (FFGs) in forested streams in the Upper Mara River basin. Source contributions are assessed in response to changes in river distance from source (RDS) as a measure of stream size (a, c, e, g, i) and density of large mammalian herbivores (LMH), b, d, f, h, j). a, b = collector-filterers, c, d = collector-gatherers, e, f = scrapers, g, h = shredders, and I, j = predators. To test the significance of the relationships, we fitted a GAM model with a smoothing function. The black line with shaded area represents smoother mean and s.e.; smoother significance, *R^2^* and GCV are supplied in the figures. Note changes on the x-axis and y-axis.
